# The formation of brain shape in human newborns

**DOI:** 10.1101/2023.01.01.521756

**Authors:** Stephan Krohn, Nina von Schwanenflug, Amy Romanello, Sofie L. Valk, Christopher R. Madan, Carsten Finke

**Affiliations:** Department of Neurology, Charité–Universitätsmedizin Berlin, Berlin, Germany; Berlin School of Mind and Brain, Humboldt-Universität zu Berlin, Berlin, Germany; Otto Hahn Group Cognitive Neurogenetics, Max Planck Institute for Human Cognitive and Brain Sciences, Leipzig, Germany; Institute of Neuroscience and Medicine (INM-7: Brain and Behavior), Research Centre Jülich, Jülich, Germany; Institute of Systems Neuroscience, Heinrich Heine University Düsseldorf, Düsseldorf, Germany; School of Psychology, University of Nottingham, Nottingham, United Kingdom

**Author notes:** Corresponding authors: Stephan Krohn, Carsten Finke, Department of Neurology Charité-Universitätsmedizin Berlin Charitéplatz 1, 10117 Berlin Germany.

**Keywords:** human brain development, human neonates, structural MRI, brain shape, topological complexity, fractal dimensionality

## Abstract

The neonatal period represents a critical phase of human brain development. During this time, the brain shows a dramatic increase in size, but it remains largely unclear how the morphology of the human brain develops in early *post-partum* life. Here we show that human newborns undergo a rapid formation of brain shape, beyond the expected growth in brain size. Using fractal analysis of structural neuroimaging data, we show that brain shape (i) strongly reflects infant maturity beyond differences in brain size, (ii) significantly outperforms brain size in predicting infant age at scan (mean error ~4 days), (iii) detects persistent alterations in prematurely born infants that are not captured by brain size, (iv) is consistently more sensitive to genetic similarity among neonates, and (v) is superior in predicting which newborns are twin siblings, with up to 97% accuracy. These findings identify the formation of brain shape as a fundamental maturational process in human brain development.

## Introduction

The human brain undergoes profound morphological changes over the lifespan (*1–3*), developing from a small and smooth structure *in utero* to the complex, highly convoluted structure that characterizes mature brains. Non-invasive studies with structural magnetic resonance imaging (MRI) have facilitated great progress in understanding these age-related changes of brain morphology, aided by the increasing availability of large open-access datasets of human MRI recordings (*4, 5*).

These advances in structural neuroimaging have recently led to the first description of normative trajectories of human brain structure over the lifespan, similar to growth charts of body weight or height (*1*). In a complementary approach, a recent line of research uses structural neuroimaging data to predict brain age from modeled trajectories of healthy brain ageing, revealing clinically meaningful discrepancies between apparent brain age and true chronological age in a variety of developmental and adult disorders (*6*).

While these advances have yielded significant insights into structural brain changes from childhood to senescence, large-scale investigations of *perinatal* brain development have remained limited, not least owing to the technical and ethical challenges of acquiring MRI data from human fetuses and newborns (*1, 3, 7*). Such investigations are vital, however, as perinatal brain maturation is fundamental for the neurotypical development of cognitive capacities and, in turn, aberrant development during this time represents a critical window of vulnerability for later cognitive deficits and neurodevelopmental disorders (*3, 8–10*).

To overcome this gap, recent collaborative efforts such as the developing Human Connectome Project (dHCP; www.developingconnectome.org) now provide the opportunity to study early-life brain development in curated datasets of unprecedented size, quality, and accessibility (*11*). These resources are met by parallel advances in the processing of early-life neuroimaging data, including neonatal brain atlases (*12–14*) and the adaptation of well-established processing pipelines to the specificities of the newborn brain regarding size variability and tissue contrasts (*15*).

Furthermore, regarding the analytical description of brain morphology, powerful new methodologies have emerged that capture the *shape* of the human brain, moving beyond the information reflected by measures of brain size, such as volume or cortical thickness. To illustrate why shape-related measures can capture additional features of brain morphology, consider the example of a fictitious structure of 10000 voxels. By definition, the volume of this structure is given by the voxels it consists of (and yields 10 ml, if voxels are 1 mm^3^ isotropic). Clearly, however, there are many ways in which these voxels could be arranged in space, resulting in different outlines of their borders or ‘shapes’ of the structure. Capturing such differences in topology, a recent line of research has shown that the shape of brain structures can be reliably described by their *fractal dimensionality* (FD) (*16–18*) – a measure of topological complexity that expresses the irregularity of a geometric shape in a single scalar number (*19–21*).

On the technical side, FD is robustly calculated from MRI segmentations of various modalities (*16, 17*), shows better test-retest reliability than volumetric measures of brain morphology (*18*), and is applicable to all tissue compartments of the brain, including cortical gray matter (GM), white matter (WM), and subcortical regions (*17, 22, 23*). This also distinguishes FD from other shape-related measures such as gyrification (*24*), which is only meaningfully applicable to the cortical sheet. Importantly, FD has proven highly sensitive not only to age-related changes of brain morphology in healthy individuals (*17, 22, 25–27*) but also to pathological alterations of brain morphology in a variety of clinical conditions including neurodegenerative, vascular, inflammatory, psychiatric, and neurodevelopmental disorders (*28–30*).

Here, we combine these recent advances of topological neuroimaging with the newly available dHCP data from newborn infants to reveal how the shape of the human brain first develops in very early life. To this end, we assess (i) the cross-sectional, longitudinal, and predictive capacity of brain shape to reflect neonatal age at the time of scanning, (ii) the impact of key neurodevelopmental factors on brain shape, including sex, singleton vs multifetal pregnancy, and premature birth, and (iii) the relationship between brain shape and genetic similarity in individual neonates. Finally, systematic comparisons between FD and volume show that brain shape complements and consistently outperforms brain size in capturing the early-life brain development of human newborns.

## Results

### Quantifying brain shape in human newborns

We analyze structural MRI recordings from the third dHCP release (*11*). This dataset includes 782 human neonates and covers a wide range of infant maturity – from very preterm to well post-term at the time of scanning (27-45 weeks post-menstrual age). Figure 1A illustrates the profound differences in brain shape over these varying degrees of maturity, where the latter are defined by the age criteria of the World Health Organization (WHO) (*31*) and the American College of Obstetricians and Gynecologists (ACOG) (*32*).

**Figure 1.**
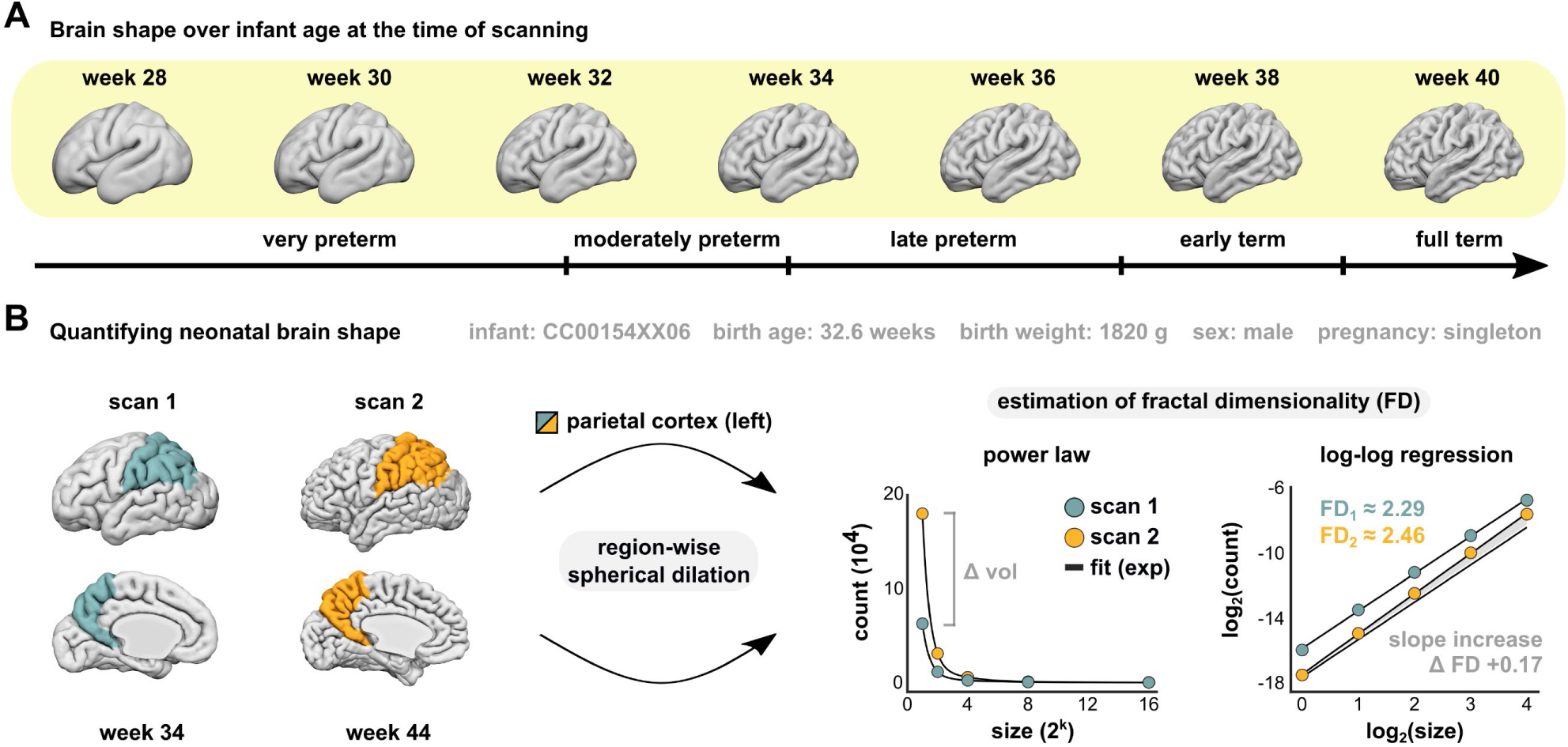
Quantifying brain shape in human newborns. **A**, Differences in brain shape over infant age at the time of scanning, illustrated for left cortical gray matter. Surface renderings correspond to the age-specific group averages of the dHCP data. The indication of infant maturity (lower arrow) follows the 2013 criteria of preterm age by the World Health Organization (*31*) and the categorization of term age by the American College of Obstetricians and Gynecologists (ACOG) (*32*). **B**, Describing the shape of a brain region by quantifying its topological complexity. The complexity estimate is calculated from spherical dilation of a region (*17, 18*), yielding its fractal dimensionality (FD) as the slope of the power law relationship between sphere size and sphere count in log-log space. This procedure measures the irregularity of a shape and is illustrated for the left parietal cortex of an exemplary infant scanned at week 34 and 44 of age. Over this ten-week interval, the change in shape (left) is reflected in an increase of topological complexity (right).

To quantify these shape differences, we apply a spherical dilation procedure to calculate the FD of each brain region, which rests on an estimation of the power law relationship between sphere size and sphere count and yields the complexity estimate as the slope of this relationship in log-log space (*16–18*). Figure 1B illustrates this procedure for the left parietal cortex of an exemplary infant scanned shortly after birth at 34 weeks and once again at 44 weeks of age. Over this ten-week interval of brain development, the change in shape that is visible from the surface renderings (left panel) is reflected by an increase in the topological complexity of that region from baseline to follow-up (right panel).

With this approach, we thus obtain one complexity estimate for every brain region (n=70) in every scan (n=884 including follow-up recordings), allowing for an in-depth evaluation of brain shape in human newborns and enabling systematic comparisons with volume as a measure of brain size.

### Brain shape reflects infant maturity beyond differences in brain size

First, we related cross-sectional age differences among newborns to the topological complexity of each brain region (measured by FD) and to the size of those regions (measured by volume). Older infants showed significantly higher topological complexity across both cortical GM and subcortical areas, with large effect sizes (Fig. 2A, left). These effects were paralleled by strong negative age-complexity correlations across widespread WM areas, such that older infants showed significantly less complex WM shapes. This GM-WM dichotomy in age-complexity correlations was further supported by an estimation of topological covariance across infants (Fig. S1), which illustrated that age-complexity effects covaried in the same direction for homologous regions across hemispheres but were strongly inversely related in GM and WM regions.

**Figure 2.**
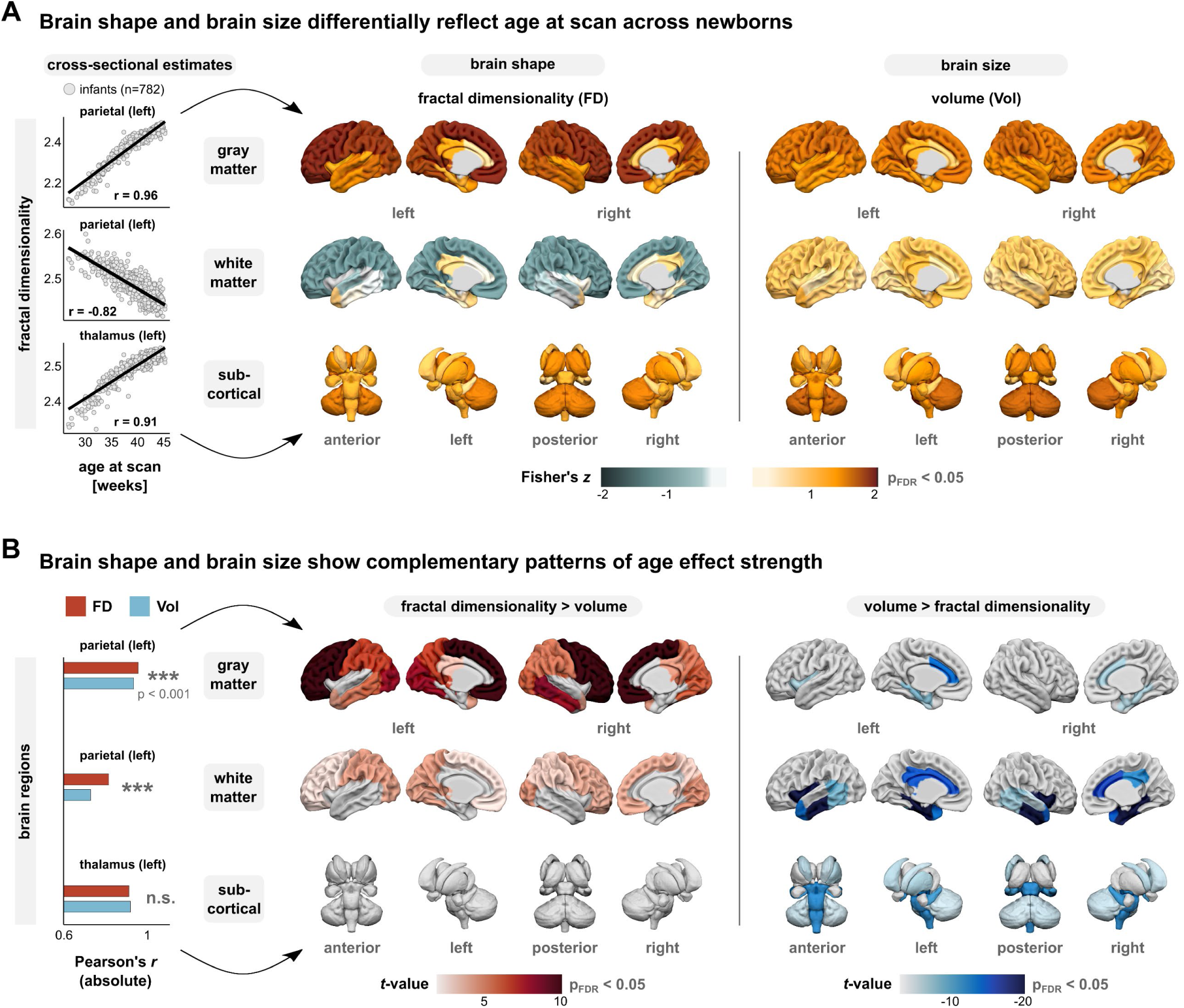
Brain shape reflects infant maturity beyond differences in brain size. **A**, Cross-sectional correlations between infant age at scan and fractal dimensionality (FD) as a measure of brain shape (left) and volume (Vol) as a measure of brain size (right). Correlation coefficients were Fisher z-transformed and thresholded to p_FDR_ < 0.05 after correction over brain regions. **B**, Region-wise comparison of age effect strength. For colored regions, the null hypothesis that fractal dimensionality and volume are equally strongly correlated with age was rejected at p_FDR_ < 0.05. Higher age correlations for brain shape are shown on the left, higher age correlations for brain size on the right. Note that in some regions such as the thalamus, infant age was reflected equally strongly by both measures.

In contrast, age-volume associations were strictly positive (Fig. 2A, right), such that brain structures were universally larger in older neonates, as would be expected from a continuous postnatal growth in brain size. While effect sizes were also large for brain volumes, a direct comparison between age-complexity and age-volume effects revealed a complementary spatial pattern, in which FD tracked infant age more strongly across most cortical GM and WM areas (Fig. 2B, left), while volume showed larger effect sizes in temporal, cingulate, and some subcortical areas (Fig. 2B, right).

Given these strong links to age, we furthermore investigated the degree to which neonatal brain shape is influenced by the sex of the infant and by pregnancy status (singleton vs multifetal). Region-wise hierarchical regression confirmed the strong age-complexity effects across the entire brain (Fig. S2A), but also revealed a significant additional impact of sex and pregnancy status on variance in FD, albeit on a much smaller scale (up to 5% additional variance explained). Notably, these effects were most pronounced in WM areas and showed spatial clusters, with infant sex primarily influencing parietal, occipital, and insular WM as well as the hippocampus (Fig. S2B), and pregnancy effects clustering in frontal, temporal, and cingulate WM (Fig. S2C).

### Longitudinal development of brain shape in individual newborns

Given these cross-sectional age-complexity associations, we next investigated how brain shape develops longitudinally *within* individual newborns. To this end, we analyzed the longitudinal trajectories of brain topology in all infants for whom repeated measurements were available (n=100). Figure 3A illustrates these individual trajectories for the occipital GM and WM of the right hemisphere. All infants showed a pronounced increase in FD from baseline to follow-up for occipital GM (paired t-test: t(99) = 25.9, p<0.001), paralleled by a simultaneous decrease in FD in the corresponding WM region (t(99)= −22.6, p<0.001; Fig. 3A), with strong effect sizes for both (Cohen’s *d* = 3.2 for occipital GM; *d* = −2.2 for occipital WM). Mapping these longitudinal developments across the whole brain revealed widespread increases in complexity across cortical GM (strongest effects in frontal and occipital lobes) as well as subcortical areas (strongest effects in the basal ganglia and thalamus), with simultaneous decreases in complexity across most WM areas (strongest effects in frontal and parietal lobes; Fig. 3B).

**Figure 3.**
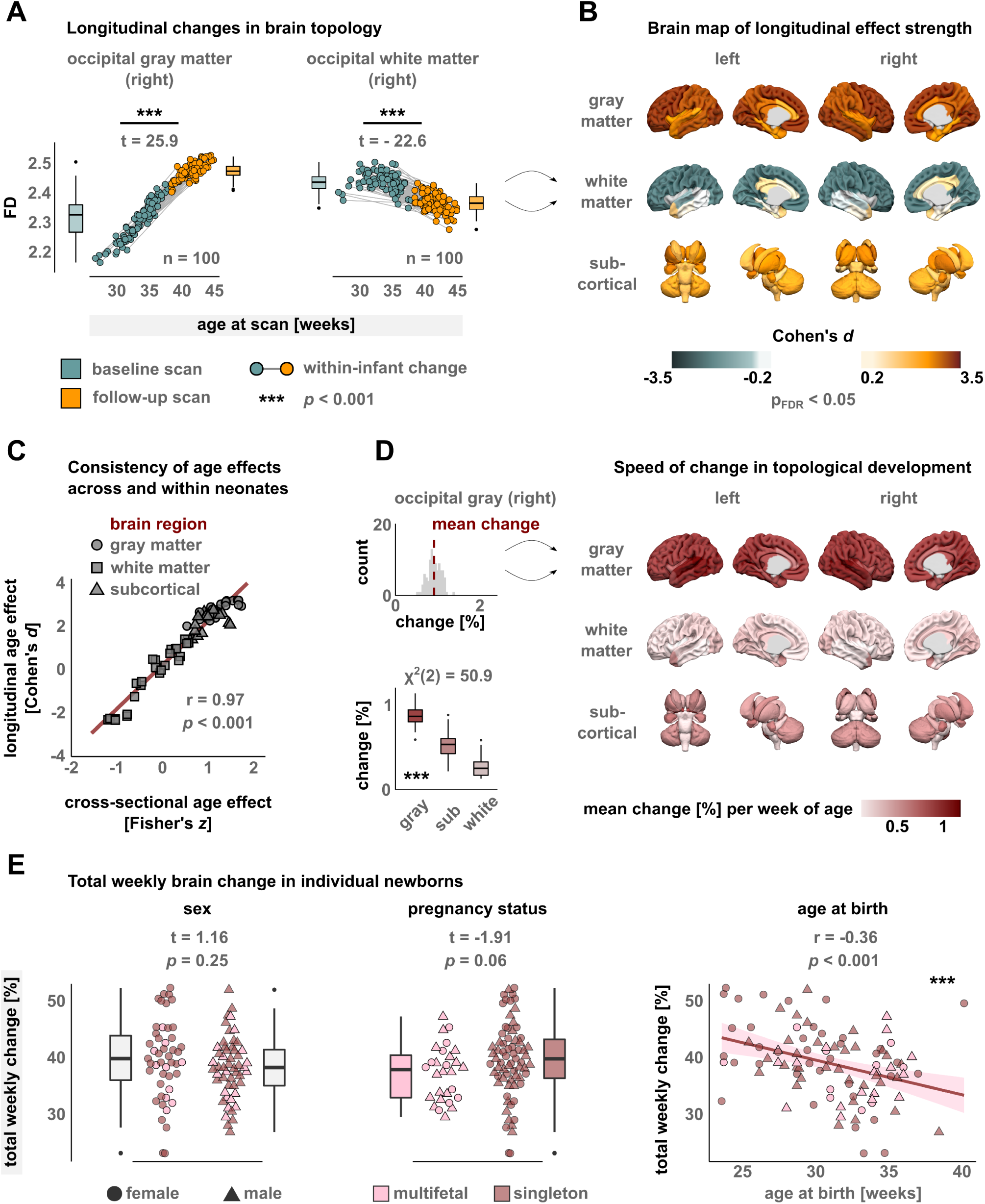
Longitudinal development of brain shape in individual newborns. **A**, Estimating longitudinal changes in brain shape. Illustration of within-infant developments for occipital GM and WM of the right hemisphere. Follow-up data were available for n=100 newborns. T-statistics derived from paired t-tests between baseline and follow-up scans. **B**, Whole-brain distribution of longitudinal age effects. Cohen’s *d* derived from the region-wise t-tests, FDR adjustment over regions. **C**, Correlation between cross-sectional age effects (Fisher’s *z*, Fig. 2) and longitudinal age effect sizes over individual brain regions. **D**, Quantifying the speed of topological development. The upper left panel illustrates the change per additional week of age for right occipital GM, where the histogram reflects individual infants. The brain map displays the mean weekly change derived from these distributions for all brain regions. The lower-left image shows the distributions of weekly change over tissue classes. Χ^2^ statistic from Kruskal-Wallis test. Pairwise comparisons between tissue classes with Dunn’s test are significant at p_FDR_<0.002. **E**, Total weekly change of brain shape in individual newborns, compared by sex, pregnancy status, and age at birth.

This spatial pattern of longitudinal age effects thus strongly paralleled the brain-wide distribution of cross-sectional age effects (Fig. 2A), including further evidence of a GM-WM dichotomy in the postnatal development of brain morphology. Indeed, explicitly comparing the distribution of cross-sectional and longitudinal estimates showed that the spatial pattern of age-complexity associations was virtually identical *across* individual newborns and *within* individual newborns (r=0.97, p<0.001; Fig. 3C).

Moreover, to characterize the spatial specificity of these longitudinal dynamics, we estimated the speed of topological development as the relative change that a brain region exhibits per additional week of age. The upper-left inset of Figure 3D illustrates this rate of change for the right occipital GM of individual infants, such that the mean of this distribution describes the average speed of development as summarized in the brain map (Fig. 3D, right). Notably, the speed of shape development showed significant differences across tissue classes (Kruskal-Wallis: χ^2^(2) = 50.9, p<0.001; lower-left inset), with cortical GM developing fastest, followed by an intermediate speed of change in subcortical areas, and WM areas showing the slowest change with age (all pairwise comparisons p_FDR_<0.002).

These dynamics furthermore raised the question of how developmental factors influence the speed of longitudinal trajectories in individual infants, such that we analyzed the total weekly brain change for each newborn (Fig. 3E). Therein, we observed no difference in the speed of development between female and male neonates (t=1.16, p=0.25), nor between singleton and multifetal pregnancies (t=-1.91, p=0.06). Interestingly, however, total weekly brain change was negatively associated with age at birth (r = −0.36, p<0.001), such that more prematurely born infants showed an accelerated development of brain topology (Fig. 3E, right).

### Brain shape outperforms brain size in predicting infant age

Given these inferential age-complexity effects, we next asked how closely infant age at the time of scanning could be *predicted* from brain shape in unseen data. To this end, we employed a supervised age prediction scheme that has been previously applied for age prediction in adults and rests on a combination of least-squares splines, dimensionality reduction, and relevance vector regression (*25, 33*). Herein, fractal dimensionality values constituted the predictor matrix, and the quality of age prediction in unseen data was assessed as the mean absolute prediction error in days (MAE) and the variance explained in the test set (R^2^), evaluated using a 10-fold cross-validation scheme (Fig. 4A).

**Figure 4.**
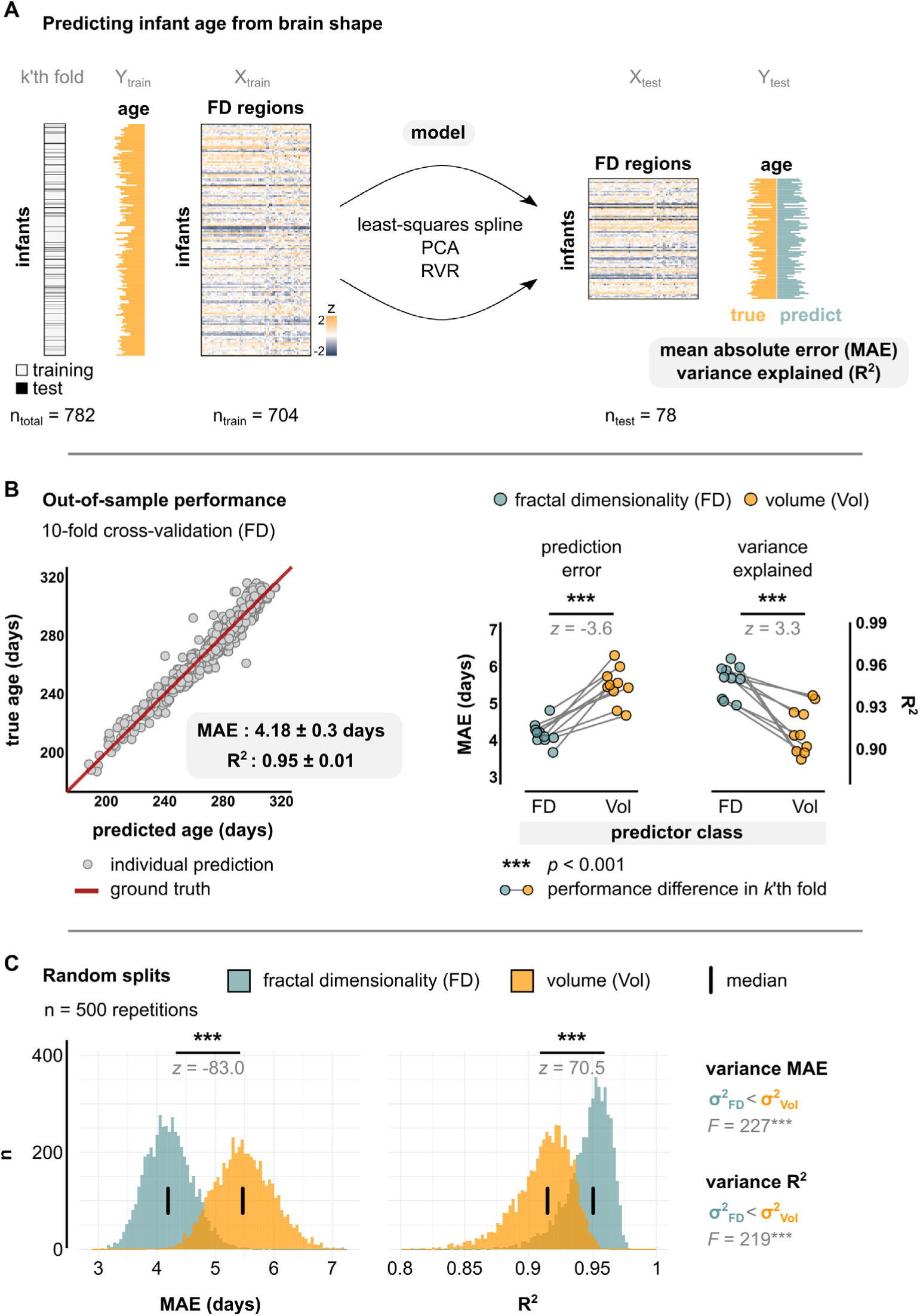
Brain shape outperforms brain size in predicting infant age. **A**, Schematic of the age prediction pipeline, resting on a combination of least-squares splines, principal component analysis (PCA) and relevance vector regression (RVR). Model performance in unseen data was evaluated by mean absolute error (MAE) of age prediction in days and variance explained in test data (R^2^), employing a 10-fold cross-validation scheme. **B**, Out-of-sample performance of predicting infant age from fractal dimensionality (left) and fold-wise comparisons between fractality-based and volume-based age prediction (right) using rank sum tests. **C**, Repetitions of cross validation procedure over random data splits to estimate the distribution of performance metrics with respect to variance and location. Differences in location between fractality- and volume-based prediction were assessed with rank sum tests, differences in variance with Levene’s test.

Out-of-sample performance of age prediction yielded very high accuracy, with a mean prediction error of 4.2 ± 0.3 days and a substantial amount of variance explained in the test data (R^2^ = 0.95 ± 0.01; Fig. 4B). Furthermore, we explicitly compared shape-based age prediction with FD to size-based age prediction with volume, showing that prediction from shape significantly outperformed prediction from size both in terms of lower prediction errors (z = −3.6, p<0.001) and more variance explained over individual folds (z = 3.3, p<0.001; Fig. 4B). Moreover, to estimate the generalizability of prediction performance over random variations in the data, we repeated the cross-validation procedure over n=500 random splits of the dataset into the ten respective folds (i.e., 5000 unique test sets) and evaluated the resulting distributions of the performance metrics for differences in location and variance. This approach not only corroborated the superior performance of FD in terms of both prediction errors (z = −83.0, p<0.001) and variance explained (z = 70.5, p<0.001), but also yielded significantly lower variance of the performance metrics for FD than for volume (MAE: F = 277, p<0.001; R^2^: F = 219, p<0.001), showing that age prediction from brain shape generalized substantially better over random fluctuations in the data (Fig. 4C).

Finally, we conducted two additional control analyses: First, age prediction from both FD and volume together performed on par with age prediction from FD alone (MAE: 4.1 ± 0.4 days, ΔMAE = 0.15 ± 0.25 days vs FD; R^2^ = 0.95 ± 0.02, ΔR^2^ = 0.5 ± 0.6 % vs FD). Second, the superior performance of age prediction from FD over age prediction from volume was equivalently observed in two alternative control models of lower model complexity (multiple linear regression and support vector regression), with virtually identical results (Fig. S2).

### Brain shape detects signatures of prematurity not captured by brain size

These pronounced relationships between infant age and brain shape furthermore raise an important general question: what normative brain shape is to be expected in infants of full-term maturity? To address this question, we estimated a reference topology at full term and quantified how much the brains of individual infants departed from this reference topology. To this end, we computed the average FD values over those infants that were both *born* and *scanned* within the full-term window, which applied to n=116 infants (Fig. 5A; full-term ACOG definition: 39 0/7 to 40 6/7 weeks; size reference was calculated in analogy from volumes). This approach subsequently allowed us to relate each individual scan to the full-term reference by computing a whole-brain index of departure from reference, based on the spatial correlation between individual topology and reference topology. Figure 5B illustrates this procedure for one infant that was born and scanned at full term and shows low departure from reference (left). Conversely, another infant that was born and scanned preterm shows higher departure from the reference topology (right). The distribution of departure indices over all scans is shown in Figure 5C and revealed that (i) departure from normative shape is significantly stronger than departure from normative size (rank sum test: z = 28.2, p<0.001), that (ii) departure indices across individual scans are significantly more variable for shape than for size (F-test: F(883,883) = 5.6, p<0.001), and that (iii) both distributions show a local minimum around term age at scan, which is expected since this is the age window on which the respective reference values were defined. Furthermore, these distributions subsequently allowed for explicit comparisons among three infant groups: (1) those born preterm and scanned preterm (*preterm-preterm*, n=161), (2) those born preterm but later scanned at term age (*preterm-term*, n=41), and (3) those born at term and scanned at term (i.e., the reference group; *term-term*, n=116). Consistent with the previously observed age effects, group 1 (*preterm-preterm*) showed significantly higher departure from the normative reference than both group 2 (*preterm-term*) and group 3 (*term-term*), and this held true for both FD and volume (Fig. 5D). In contrast, the comparison between group 2 (*preterm-term*) and group 3 (*term-term*) was only significant for FD, but not for volume (Fig. 5D), showing that brain shape captured persistent alterations in prematurely born infants, even when those infants were later scanned in the full-term age window, while such signatures of prematurity were not detected with brain size.

**Figure 5.**
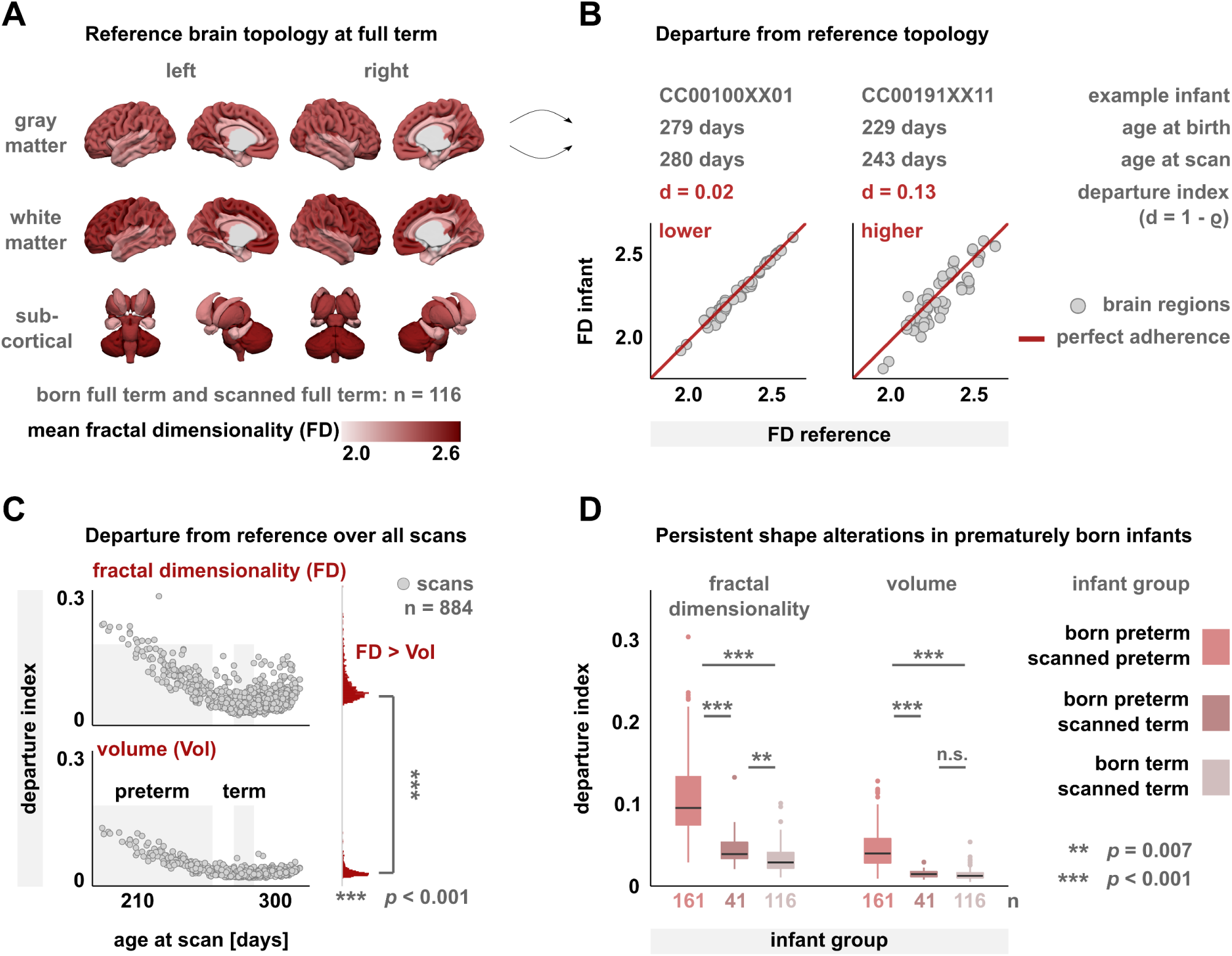
Brain shape captures persistent alterations in prematurely born infants that are not detected by brain size. **A**, Reference topology derived as the mean fractal dimensionality (FD) per region over all infants that were both born and scanned at full term (n=116). Term is defined as 39 0/7 weeks to 40 6/7 weeks of age based on the ACOG criteria (*32*). **B**, Quantifying the departure from this reference topology with a departure index *d*, computed from the spatial rank correlation between each infant’s individual topology and the reference values of panel A. Illustration for two infants with a lower-departure scan (left; born term, scanned term) and a higher-departure scan (right; born preterm, scanned preterm). Reference brain size was computed in analogy using regional volumes. **C**, Departure from reference over all n=884 scans in the dataset for brain shape (top) and brain size (bottom). The shaded areas display the ACOG definitions of preterm age (< 37 0/7 weeks = 259 days) and term age (273 to 286 days). Note the local minimum of both scatter clouds around the term window. Departure indices were significantly higher for FD than for volume (p<0.001, rank-sum test). **D**, Departure from reference for three infant groups: (1) born preterm and scanned preterm, (2) born preterm and scanned term, and (3) born term and scanned term. Kruskal-Wallis omnibus tests yielded significant results for both FD (χ^2^(2) = 197.2, p<0.001) and volume (χ^2^(2) = 194.6, p<0.001). Pairwise comparisons correspond to Dunn’s tests with FDR adjustment.

### Brain shape reflects genetic similarity among individual newborns

Having shown a principled link between infant age and brain shape on the group level, we lastly sought to move beyond age effects and study the relationship between genetic factors and the variability of brain shape on the level of individual newborns.

To this end, we computed the pairwise age differences for all infant-to-infant comparisons in the dataset and measured the topological dissimilarity of any two children (i.e., the ‘shape difference’) as the distance between their whole-brain fractality profiles (Fig. 6A). As expected from the group-level age effects, the shape difference between any two infants strongly increased with the age difference between them (ρ=0.83, p<0.001; Fig. 6B). However, the granularity of individual brain-to-brain comparisons subsequently allowed us to threshold the pairwise age difference to obtain only those comparisons in which both infants were within 1 day of age at the time of scanning. The inset of Figure 6B shows that, even within this subset of age-matched comparisons, there is considerable variance in the dissimilarity of individual brain topologies. Importantly, however, these shape differences are not attributable to age because the respective infants were the same age at the time of scanning, allowing us to evaluate if sharing genetic information —beyond sharing the same age—would be linked to a higher similarity in brain shape.

**Figure 6.**
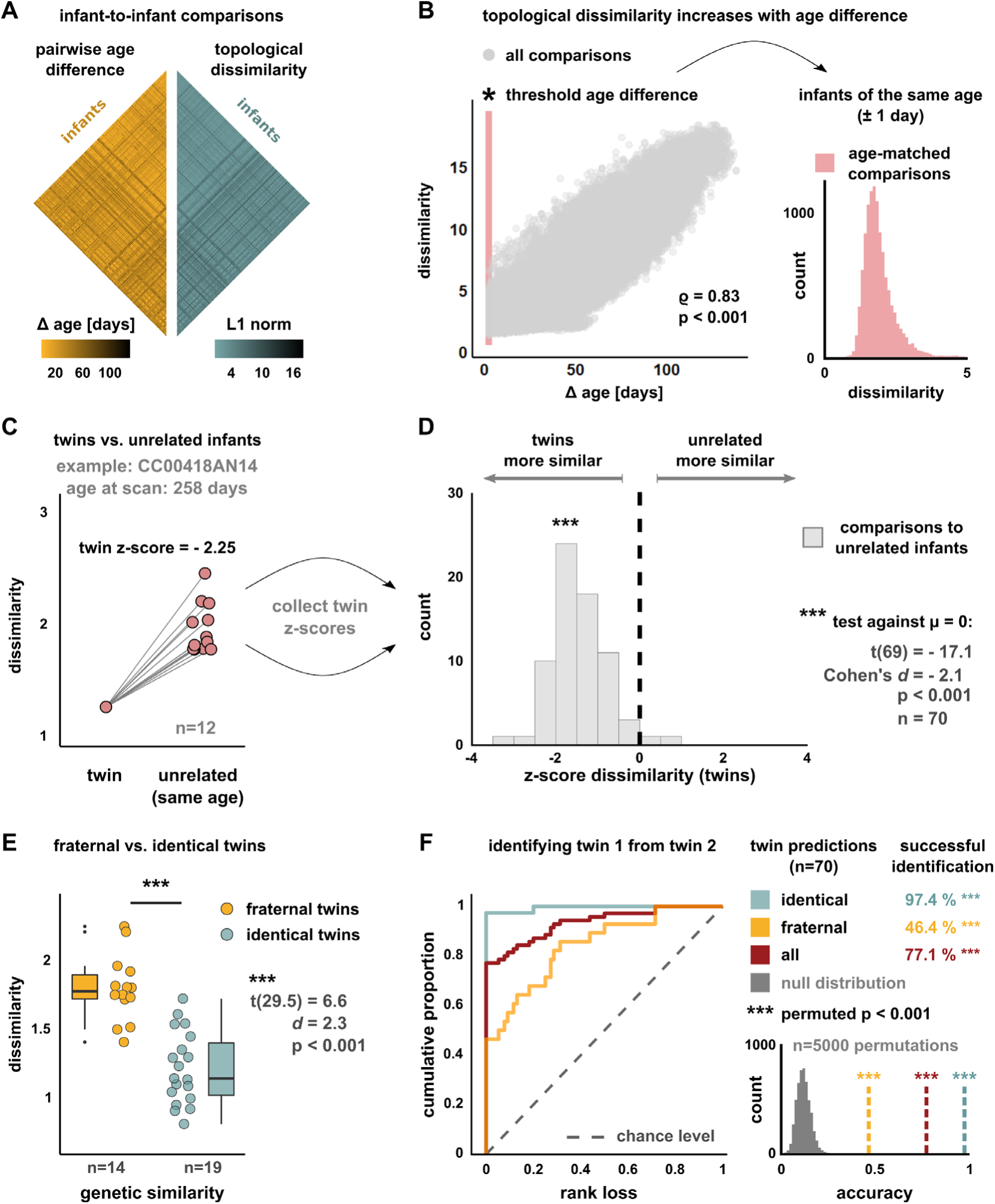
Brain shape reflects genetic similarity among newborns and enables the prediction of one twin from the brain of the other twin (‘twingerprinting’). **A**, Pairwise age differences and topological dissimilarity (‘shape difference’) between any two infant brains in the dataset. **B**, Correlation between age difference and topological dissimilarity for all infant-to-infant comparisons. Age-matched dissimilarity distribution after thresholding the age difference to ±1 day. **C**, Dissimilarity scores between an exemplary infant and its twin sibling (left) and all unrelated infants of the same age (right). The dissimilarity of the twin sibling was z-scored with regard to all age matches and collected for each twin-to-unrelated comparison (n=70). **D**, Distribution of the twin dissimilarites from panel C over all twin-to-unrelated comparisons. One-sample t-test against zero. **e**, Dissimilarity scores across fraternal and identical twin pairs (Welch’s t-test). **F**, Identification of twin siblings from unrelated infants of the same age. For each twin-to-unrelated comparison (see panel C), the infant with the lowest-ranking disimilarity was predicted to be the target twin. The ROC-like curve shows the proportion of infants over increasing rank loss (0: correct identification; 1: all unrelated infants more similar than target twin). The null distribution of twin predictions was estimated by randomly permuting the rank structure and recording the correct twin identifications that happen by chance.

To test this idea, we first compared the brains of twin siblings to all matched infants that were the same age as these twins but were biologically unrelated to them. Figure 6C illustrates the resulting dissimilarity distribution for one of the 35 twin pairs for whom unrelated matches were available. Herein, the difference between the exemplary infant and its twin sibling was substantially lower than the difference to any of the unrelated children, such that the two twin brains were the most alike in shape within this age-matched group. Critically, this observation generalized over all twin-to-unrelated comparisons — brain shapes of twin siblings were generally more similar to each other than to the brains of unrelated infants, with large effect size (one sample t-test: t(69) = −17.1, p<0.001, Cohen’s *d* = −2.1; Fig. 6D).

As this finding strongly suggested a link between brain shape and genetic similarity, we performed two additional analyses to test the idea that similarity in brain shape may reflect similarity in genetic information.

First, we stratified the dissimilarity scores by the sex of the compared infants (Fig. S4). This revealed that infants of the same sex exhibit significantly more similar brain shapes than infants of different sexes, and this was true both in twin siblings and in biologically unrelated infants. Interestingly, for infants of the same sex, brain topologies were even more similar when both newborns were female compared to when both newborns were male (z = −6.2, p_FDR_<0.001 for unrelated, tendency in twins; Fig. S4), suggesting an additional effect of homologuous sex chromosomes that share the same genes (i.e., an XX karyotype in both infants) compared to heterologuous sex chromosomes (i.e., an XY karyotype) that do not. Second, we hypothesized that, even among individual twin pairs, sharing more genetic information would be expressed in more similar topologies still. Accordingly, we stratified twin pairs into dizygotic siblings (i.e., fraternal twins with ~50% shared genes) and monozygotic siblings (i.e., identical twins with ~100% shared genes). Critically, we indeed observed that brain shapes are significantly more similar in identical twins than in fraternal twins (t(29.5) = 6.6, p<0.001, d = 2.3; Fig. 6E).

Notably, analogous control analyses with volume again showed that genetically related infants exhibit much stronger similarity in brain shape than in brain size (Fig. S5).

### *Twin*gerprinting – identifying the brain of one twin from the brain of the other twin

Given these links between brain topology and genetic similariy, we lastly asked if the similarity of brain shape would enable the prediction of twin siblings out of the set of matched unrelated infants (Fig. 6F). This approach pertains to the idea of ‘connectome fingerprinting’ (*34*), in which the unique variability of brain activity signatures (‘fingerprints’) enables the identification of single individuals out of large samples with high accuracy. Importantly, however, here we do not aim to identify the same individual, but its twin silbing (‘*twin*gerprinting’). To this end, the dissimilarity scores of individual twin-to-unrelated comparisons were ranked, and the infant with the lowest-ranking shape difference was predicted to be the twin sibling of the target infant (i.e., the other twin). In the example of Figure 6C, the twin sibling was thus correctly identified, but not so in the analogous analysis with volume (Fig. S5). To assess the predictive power of this approach, three metrics were evaluated: (i) we computed the ‘rank loss’ over individual predictions, defined as the proportion of unrelated infants whose brain shapes were more similar to the target infant than its twin (i.e., rank loss = 0: correct identification; rank loss = 1: all unrelated more similar than twin; Fig. 6F, left), (ii) we computed the accuracy of twin predictions as the proportion of correction identifications, and (iii) we estimated the null distribution of correct twin identifications that happen by chance. The latter was implemented by randomly permuting the ranks within individual predictions, yielding the permuted p-value on the prediction accuracy as the proportion of randomly obtained accuracies that surpass the empircially observed value (lower-right inset). On average, ~11% of twin identifications are thus expected to happen by chance.

Critically, Figure 6F shows that brain shape correctly identified the target twin in 77.1% over all predictions (p_perm_<0.001). Notably, however, predictive power again mirrored the effect of genetic similarity on brain shape: while the accuracy of identifying fraternal twins was considerably lower (46.4%), if still far from chance (p_perm_<0.001), prediction accuracy was near-perfect in the case of identical twins (97.4%, p_perm_<0.001; Fig. 6F).

Finally, analogous analyses with volume again showed that predictive power of brain size was markedly lower, resulting in a consistent 25-30% drop in identification accuracy and approaching chance levels in the case of fraternal twins (Fig. S6).

## Discussion

These findings show that human newborns undergo a rapid formation of brain *shape*, beyond the expected growth in brain *size*. By providing a principled account of the brain’s morphological changes over the first few weeks after birth, we show that this early-life formation of brain shape represents a fundamental maturational process in human brain development.

To this end, we analyze structural neuroimaging data from the developing Human Connectome Project – one of the largest datasets of human newborns ever collected (*11*). Therein, we describe neonatal brain shape with *fractal dimensionality* —a geometric measure of topological complexity which complemented and systematically outperformed purely size-based accounts of early-life brain development. In particular, we find that (i) brain shape strongly reflects infant maturity beyond size differences, both cross-sectionally and longitudinally, (ii) brain shape consistently outperforms brain size in predicting infant age in unseen data, with high accuracy (mean error ~4 days), (iii) brain shape detects persistent alterations in prematurely born infants that are not captured by brain size, (iv) brain shape is consistently more sensitive to genetic similarity among newborns, assessed by comparing infant sex, related vs unrelated infants, and fraternal vs identical twins, and (v) brain shape enables the identification of one twin from the brain of the other twin (‘*twin*gerprinting’) with high accuracy (~77% overall, ~97% in identical twins), again outperforming twin predictions from brain size.

Below, we turn to the implications of these findings, which advance our understanding of early-life brain development in five key directions.

First, brain shape is inextricably linked to infant age. The topological complexity of neonatal brains strongly reflected age at the time of scanning, capturing differences in infant maturity based on international consensus criteria. Notably, this held true both for developmental differences *across* individual newborns (i.e., cross-sectional variation) and for developmental differences *within* individual newborns (i.e., longitudinal variation, as individual infants mature). Therein, the direction of shape development was highly consistent across both, showing a dichotomy between cortical GM that grows increasingly more complex with brain maturation and WM regions that grow increasingly less complex as the brain matures. In contrast, brain volumes showed strictly positive associations with age, as would be expected from a continuous postnatal growth in brain size. Notably, the inverse age-complexity associations in WM were strong, wide-spread, and ubiquitously observed across individual newborns. This increasing WM regularity is thus likely to reflect early-life myelination, which is thought to start in the second half of pregnancy and last well into adolescence (*7*). Given that higher myelin content shifts voxel intensity gradients to a more WM-like spectrum, lower WM complexity in more mature newborns may thus be an expression of increasingly smoother WM boundaries with ongoing myelination. Moreover, this spatial dichotomy in GM and WM development was paralleled by a temporal dichotomy, in which cortical GM showed the most rapid change over time, while WM topology showed a much more protracted evolution.

These findings in neonates align remarkably well with pioneering work on brain growth trajectories over the first two years of life, which reported a much slower development of WM compared to cortical GM (*35*). Here we not only corroborate the different speeds of development in GM and WM, but also show that such tissue-specific dynamics are already present right after birth. Notably, these perinatal dynamics also converge with a recent account of normative brain growth over the larger lifespan (*1*), which suggested that developmental trajectories may be steeper for GM than for WM around birth.

Second, it is particularly worth focusing on the early-life development of cortical topology, which constituted some of the strongest effects throughout our study. Overall, our results suggest that the dynamic increases of topological complexity in the cortex are an expression of early-life cortical folding. This folding process accelerates markedly around 26 weeks of gestational age, when the brain begins a rapid change from a near-lissencephalic to a highly convoluted structure *in utero* (*36–38*). Here we show that this morphological development naturally extends into the neonatal period, where the increasing cortical convolution (Fig. 1) becomes apparent as a highly canonical increase in topological complexity (Fig. 2–3). Interestingly, recent evidence from statistical physics suggests that the cortical topologies observed across a variety of primate species may be an expression of the same archetypal fractal shape (*39*). Given the highly canonical shape developments observed here, this raises the intriguing possibility that the early-life formation of cortical complexity is not only a key process in human brain development but may rather be the result of a more general, evolutionarily conserved mechanism of cortical expansion.

Third, we show that differences in age do not only *explain* differences in brain shape, but that this relationship can be inverted to *predict* the age of an infant from the shape of its brain with high accuracy (mean error ~4 days). Here again, brain shape significantly outperformed brain size, and this was consistently observed across performance metrics, data splits, and three different prediction models. Notably, this high accuracy was homogeneously observed across the whole age range in the dataset, from very premature (~28 weeks) to well after term (~44 weeks). Together with the group-level age effects, this shows that brain shape closely reflects infant maturity over all stages of neonatal development and identifies shape development as a highly canonical process of early-life brain maturation.

Fourth, we show that brain shape captures persistent morphological alterations in prematurely born infants that remained undetected by brain size. Specifically, even when premature infants were subsequently scanned in the full-term age window, their brain topologies still deviated significantly from a reference topology of term-born infants, whereas this was not the case for a normative reference of brain size. While these findings reveal a persistent developmental lag in the spatial organization of brain morphology, we additionally observed significant differences in the temporal trajectories of preterm infants, where the most prematurely born children exhibited an accelerated development of brain topology. These findings show that brain shape reflects altered developmental trajectories of preterm infants already at a very early stage of postnatal life. While this is —to the best of our knowledge— the earliest account of altered topological dynamics in human brain development, one previous study applied fractal analysis to global GM and WM segmentations in infants at 12 months and found that prematurely born infants with intrauterine growth restriction showed persistent reductions in GM complexity that were related to language and motor scores (*40*). Moreover, a recent study has reported persistent reductions of cortical complexity at adult age in those participants that had been born prematurely: these alterations clustered in temporoparietal areas, were related to the severity of prematurity at birth, and correlated with reduced cognitive performance in adulthood (*41*). These findings not only align well with the topological alterations that we observe here early on in preterm neonates, but also suggest that these changes may carry important functional significance for the neurocognitive development in later life.

Importantly, about 11% of infants are born prematurely word-wide (*31*), bearing an increased risk for early-life mortality (*31, 42*), later-life cognitive deficits (*8*), and neuropsychiatric disorders (*43*). This risk profile has motivated the application of advanced neuroimaging techniques in search for prognostic biomarkers of neurodevelopmental outcomes after premature birth (*10*). Although large-scale follow-up will be necessary to evaluate the prognostic value of shape alterations systematically, our findings suggest that brain topology measures such as fractal dimensionality represent highly promising candidates for early-life imaging markers of at-risk neurocognitive development. Therefore, long-term longitudinal efforts are urgently needed to follow up neonates into infancy and adulthood when neurodevelopmental disorders are manifest.

Fifth, our study reveals a fundamental link between neonatal brain shape and genetic information. By focusing on the morphological variability of individual brains, we show that the degree to which any two brains are similar in shape is strongly associated to the genetic similarity of the compared infants. Specifically, we find that (i) the brains of genetically related infants are more similar in shape than those of unrelated infants, (ii) infants of the same sex show more similar brain shapes than infants of different sexes, (iii) brain shapes are more similar in homologous sex chromosomes than in heterologuous sex chromosomes, and that (iv) brain shapes are more similar in identical twins (~100% shared genes) than in fraternal twins (~50% shared genes). Importantly, all these comparisons were carried out in age-matched infants, such that these results are unlikely to be confounded by the strong age effects discussed above.

These results provide a critical addition to the fast-growing literature linking neuroimaging phenotypes to genetic factors in human brain development (*2, 44–48*). In this regard, one recent study has shown that cortical morphology at birth reflects spatiotemporal patterns of gene expression in the fetal human brain (*49*), suggesting that the topological maturation observed here *post-partum* is a direct extension of intrauterine genetic regulation. Moreover, the impact of infant sex on neonatal shape similarity we observe here aligns well with recent reports of greater variability of brain structure in males than in females over the larger lifespan (*50–52*). Furthermore, a recent study found that deviations from normative brain age in adulthood were best explained by congenital factors such as polygenetic risk, suggesting that early-life genetic factors exert a lifelong influence on brain structure (*53*).

Finally, the strong links between genetic information and brain shape enabled us to predict which infants are twin siblings by identifying the brain of one twin from the brain of the other twin. The idea of identifying individuals from neuroimaging data pertains to the approach of ‘connectome fingerprinting’ (*34, 54*), in which unique patterns of brain activity (‘fingerprints’) allow for the identification of single individuals with high accuracy. Importantly, however, here we did not aim to identify the same individual, but its twin silbing (‘*twin*gerprinting’). Indeed, we were able to identify twin siblings from the similarity of their brain shapes with high accuracy (~77% overall, ~97% in identical twins), and here again, brain size was much less sensitive in detecting genetic similarity among newborns. Overall, these findings suggest that shape similarity is a direct expression of genetic similarity, and that the variability of individual brain shapes represents a genetically modulated and heritable phenotype in human newborns.

In sum, our study identifies the early-life formation of brain shape as a fundamental maturational process in human newborns, with several immediate implications for our understanding of normative brain development, the study of neurodevelopmental disorders, and the relationship between morphological variability and genetics.

## Methods

### Data and image processing

All data analyzed here were obtained from the third release of the developing Human Connectome Project in 2021 (dHCP; www.developingconnectome.org), including cross-sectional data for n=782 infants (360 females, 422 males). Magnetic resonance imaging of virtually all newborns was acquired during natural sleep (*11*). Mean birth age was 37.89 ± 4.17 weeks [range: 23.0 – 43.57], and age at first scan was 39.81 ± 3.55 weeks [range: 26.71 – 45.14]. Of these infants, 682 were born from singleton pregnancies, while 100 were born from multifetal pregnancies. Follow-up scans for longitudinal analyses were available for n=100 infants. Note that compared to adult brains, tissue contrasts in neonatal brains are inverted due to immature myelination (*3, 55*), such that T2-weighted images provide better quality and were hence used for image processing in the dHCP (*15*), including surface reconstruction with FreeSurfer (*56*).

Brain segmentations of individual images were provided with the dHCP data and were based on the Draw-EM algorithm (Developing brain Region Annotation With Expectation-Maximization) (*13, 15*). Therein, assignment of individual voxels (0.5 mm^3^ isotropic) to regions of interest (ROI) rests on the ALBERT atlas for neonatal brain anatomy (*12*). This atlas contains 87 regions, including 16 cortical gray matter and white matter regions for each hemisphere, 9 bilateral subcortical regions, the brainstem and corpus callosum as unpaired regions as well as unlabeled tissue, background, and cerebrospinal fluid. For topological analysis, we discarded these latter labels and furthermore combined some smaller and contiguous regions to harmonize spatial granularity across the brain. Specifically, we combined the medial and lateral part of the anterior temporal lobe, the anterior and posterior segments of the gyri parahippocampalis et ambiens, the anterior and posterior lateral occipitotemporal gyrus as well as high-intensity and low-intensity voxels of the thalamus, yielding a total of 70 ROIs assigned in each MRI.

Notably, the dHCP provides age-specific normative templates by week of post-menstrual age to account for the rapid development of neonatal brains. These age-specific group-averages are openly available from https://brain-development.org (*14, 57, 58*), and the surface renderings of the left cortical gray matter correspond to the week-wise averages displayed in Figure 1A. For all visualizations of statistical tests, results were mapped onto the 40-week template (Fig. 2,3,5, and S3).

### Estimating fractal dimensionality

As summarized in Figure 1, the shape of a brain region was quantified by its fractal dimensionality (FD), using the openly available calcFD toolbox (https://github.com/cMadan/calcFD) for MATLAB (The MathWorks Inc., Natick, Massachusetts). In general, FD can be measured empirically by estimating the scaling law between the size of a measurement unit and the number of those units required to cover the object comprehensively in an embedding dimension (here: 3D voxel space). To estimate FD from the structural MRI segmentations, we here apply the spherical dilation algorithm from the calcFD toolbox (*17, 22*), which is computationally similar to the traditional box-counting method (*16*), but is less sensitive to object translation and rotation (*17*) and yields better test-retest reliability than box-counting (*18*). Computationally, empirical power law estimation requires a definition of the physical scales over which the estimate is computed (*16*), i.e., the range of voxel sizes, typically expressed as 2*^k^* with *k* ∈ ℕ_0_. Here we follow previous applications of the toolbox in applying the canonical range of *k* = 0,1, …,4 for FD estimation with spherical dilation (*17, 18, 22, 25*). Finally, we extended the calcFD toolbox to process the neonatal brain parcellations described above. In brief, all voxels assigned to a particular region were indexed, yielding a binary 3D mask of that region which was subsequently passed to spherical dilation, as illustrated for the left parietal cortex in Figure 1B. This process was iterated over all parcels to obtain the FD value of each region, yielding a 1 × 70 vector for each scan. As the parcellated images contain isotropic voxels (*15*), the volume estimate of a region was computed as the sum of all voxels assigned to that region by the segmentation.

### Inferential statistics and modelling

All directional tests were two-tailed. Simple two-group comparisons were tested with t-tests or rank sum tests, depending on the distribution of the variables, and in analogy for correlational analyses with either product-moment or Spearman’s rank correlation. Two-sample tests were unpaired except for the longitudinal analyses in Figure 3A, in which each newborn had a baseline and a follow-up scan, such that these distributions were considered paired samples. Effect sizes for parametric group tests were computed as Cohen’s *d*. Parametric correlation strengths were Fisher r-to-z-transformed to harmonize scales for the visualizations in Figures 1A and S1. Multiple-group omnibus tests (Fig. 3D, 5D, S4) were implemented with Kruskal-Wallis tests, followed up by pairwise Dunn’s tests. Formal significance was considered at an α-level of 0.05, and p-values of multiple pairwise tests were corrected after Benjamini-Hochberg (*59*) to control the false discovery rate (FDR). For the comparison of correlation coefficients in Figure 2B, the null hypothesis posits that two variables (FD and volume) are equally correlated with a third variable (age at scan), all obtained from the same individuals (*60*). Furthermore, for the topology covariance network in Figure S1, the pairwise region-to-region correlation matrix of FD values was constructed from the cross-sectional scans in Figure 2, and this matrix was thresholded to the top and bottom first percentile to obtain the strongest positive and inverse covariance across brain regions. Finally, the hierarchical regression in Figure S2 compared a compact model in which the fractal dimensionality of a brain region was explained with infant age alone (FD ~ age) to two augmented models which incorporated sex (FD ~ age + sex) and pregnancy status (FD ~ age + pregnancy), respectively. To estimate in which brain regions these factors significantly explained additional variance beyond age, compact and augmented models were compared with F-tests for nested models, using the lmSupport package for R (https://rdrr.io/cran/lmSupport).

### Predicting infant age

The age prediction pipeline in Figure 4 rests on the openly available PRISM toolbox (https://github.com/cMadan/prism) for MATLAB, which was developed for age prediction from brain features and includes a combination of least-squares splines, dimensionality reduction, and relevance vector regression (*25, 33*). Herein, the smoothing parameter for spline regression was set to zero, enforcing near least-squares cubic spline to counteract overfitting; all other parameters were left to default, including the application of principal component analysis and relevance vector regression within a sparse Bayesian framework (*61*). The predictor matrix was of the form [observations × brain features] and contained either fractal dimensionality values, volumes, or both. All predictors were standardized. To evaluate prediction performance, we applied a standard 10-fold cross-validation scheme, such that the model was trained on ~90% of the data and predicted age at scan in the remaining 10% of the data in each iteration. Note that we here limited the dataset to the 782 unique baseline scans (i.e., excluding the follow-up sessions) to ensure that every infant contributed exactly one scan to the data. Prediction quality for each iteration was then assessed as the mean absolute prediction error (|predicted age - true age|) and the variance explained in the test set (1 – residual sum of squares / total sum of squares), as shown in Figure 4B. For the random repetitions of the cross-validation procedure (Fig. 4C), we computed 500 unique permutations of the data that were subsequently split into ten folds, resulting in 5000 predictions on unique test sets. Finally, to assess the impact of different model types, we applied the same prediction pipeline using simple multiple linear regression and Support Vector Regression with a linear kernel with MATLAB-inbuilt functions (*fitlm* and *fitrsvm*), as shown in Figure S3.

### Departure from normative reference

For the analyses in Figure 5, we estimated reference values of brain shape and size in infants of full-term maturity. This approach is conceptually related to the hub disruption index (*62*) in functional neuroimaging, in that data points from single individuals are compared to normative data points obtained from a reference population. Here, the reference population consisted of those infants that were both born and scanned within the full-term window, where the latter was defined based on the ACOG definitions (39 0/7 weeks to 40 6/7 weeks). This criterion was fulfilled by n=116 newborns in the dataset. For each brain region, the full-term reference value was then computed as the average over those 116 infants, once for FD values (shape reference; Fig. 5A) and once for volumes (size reference). This approach subsequently allowed for a comparison between the reference values across all brain regions and the corresponding values computed from individual scans, as shown in the scatter plots of Figure 5B. To estimate how much these individual scans deviated from the full-term reference, we computed a departure index defined as d = 1 − ϱ, that is, the nonparametric spatial correlation distance between the individual scan and the reference. Therein, Spearman’s rank correlation was chosen because (i) we aimed to obtain an estimate of the relative spatial organization across the whole brain and because (ii) the speed of development varied over the different tissue classes (Fig. 3D), such that the deviations from reference were not uniform but showed clustering effects (e.g., deviations cluster below the identity line in Fig. 3B). For each scan, we thus obtain one index of departure from full-term shape reference (FD) and another for the departure from full-term size reference (volume). These indices were subsequently compared across all scans (Fig. 5C) and among infants that were born preterm and scanned preterm and those that were born preterm but scanned later at term-equivalent age based on the ACOG definitions (Fig. 5D). Finally, note that infants who met the full-term criterion are expected to follow the reference closely because they formed part of the group on which this reference was defined, thus providing an estimate of variability within the full-term group itself. This close adherence to reference was indeed observed for both FD and volume in full-term infants (Fig. 5B-D).

### Comparing individual infant brains

Apart from the above group-level inferences, we furthermore conducted comprehensive pairwise comparisons of individual neonatal brains (Fig. 6). For any two given infants, we thus quantified the overall ‘shape difference’ of their brains by taking the vectors of their regional FD values and computing the dissimilarity between these two vectors. To this end, we here apply the L1 norm (‘Manhattan distance’), as this measure weights all vector components equally and is less sensitive to single-dimension deviations compared to the Euclidean distance, since the individual terms are left unsquared. For every infant-to-infant comparison, this approach yields a scalar measure of overall dissimilarity (Fig. 6A), such that higher values indicate more pronounced shape differences and lower values indicate that the compared brains are more similar in shape. Moreover, the identical approach was applied to regional volumes to compute the overall dissimilarity in size between any two brains (Fig. S5).

### Genetic similarity

These brain-to-brain comparisons subsequently allowed us to relate the shape similarity of any two brains to the genetic similarity of the compared infants. The latter was formalized in three different sets of comparisons: (i) infants of the same sex vs infants of different sexes (Fig. S4), (ii) twin siblings vs unrelated infants (Fig. 6C-D), and (iii) identical twins vs fraternal twins (Fig. 6E-F). Overall, there were 42 twin pairs in the dataset. For the age-matched analyses in Figure 6, however, a total of seven twin pairs had to be discarded, one because no unrelated infants of the same age were available, and six because the two twin siblings themselves were scanned more than one day apart, leaving 35 twin pairs. Moreover, the genetic similarity among those twin pairs was further assessed by categorizing them into identical twins (i.e., monozygotic siblings) and fraternal twins (i.e., dizygotic siblings). This information on twin status was provided by the dHCP consortium (Dr Harriet Cullen, King’s College London) and was derived from single nucleotide polymorphisms (SNP) array genotype data, which were used to confirm whether the twins were monozygotic, sharing 100% of their genetic variation (PI_HAT = 1), or dizygotic, sharing approximately 50% of their genetic variation (PI_HAT ~ 0.5) (*63*). These data on twin sibling status were available for 33 twin pairs.

### Twin predictions

Apart from the inferential effects of sex and twin status reported in Figure 6C-E and Figure S4, we furthermore predicted twin siblings out of the set of age-matched unrelated infants (Fig. 6F). To this end, we iterated over all individual twins-to-unrelated comparisons and predicted the lowest-ranking dissimilarity score (i.e., the most similar brain in shape) to belong to the twin of the target infant, as detailed in the main text and Figure 6C. Note that although the set of unrelated matches was the same for any two twins, the dissimilarity scores between twin A and the unrelated infants and twin B and the unrelated infants naturally differed, as all these comparisons were individual pairwise measures. In consequence, every twin pair resulted in two predictions –once identifying twin B from twin A and once identifying twin A from twin B— yielding 70 twin predictions in total. Furthermore, note that the number *n* of unrelated matches varied across the individual twin pairs, such that the chance level of individual twin predictions varied in parallel as 1⁄*n*. For illustration, the example of Figure 6C features 13 age-matched infants (of whom one was the twin to be identified), resulting in a chance level of 1⁄13 ≈ 7.7 %. As such, chance levels for individual predictions were higher if fewer unrelated matches were available in the dataset (maximum 50% if only one unrelated match was present). To account for this variability, we implemented a permutation approach in which the rank structure within individual predictions was randomly shuffled 5000 times and the proportion of chance identifications was recorded over all individual predictions. In consequence, we obtain a null distribution of correct twin identifications that happen by chance, which yields the p-value of the empirically observed identification accuracy was the proportion of permuted accuracies that surpass the empirical value. The inset of Figure 6F shows this null distribution, which yielded a mean accuracy of 11.4 ± 3.7 % of correct twin identifications that happen by chance.

Finally, the identical approach was applied to twin prediction from brain volumes, allowing for an explicit comparison between shape-based and size-based twin prediction (Fig. S6).

## Acknowledgments

We would like to thank Dr Harriet Cullen at King’s College London for providing us with the information of genetic relationships among twin siblings (dizygotic vs. monozygotic). SK was funded by the German Research Foundation (DFG) grant number FI 2309/2-1. SLV was supported by Max Planck Gesellschaft (Otto Hahn award) and the Helmholtz International BigBrain Analytics and Learning Laboratory (HIBALL). CF was supported by German Research Foundation (DFG) grant numbers 327654276, FI 2309/1-1 and FI 2309/2-1 and the German Ministry of Education and Research (BMBF), grant numbers 01GM1908D, 01GM2208C and 01GM2102.

## Data and code availability

All data analyzed here are openly available from the developing Human Connectome Project (http://www.developingconnectome.org). Preprocessed data as well as analysis code are available from the corresponding author and will be made publicly available on the Open Science Framework upon acceptance of the manuscript (https://osf.io/6jck4/).

## Supplementary information

Supplementary Figures S1-6

---

**Figure S1.**
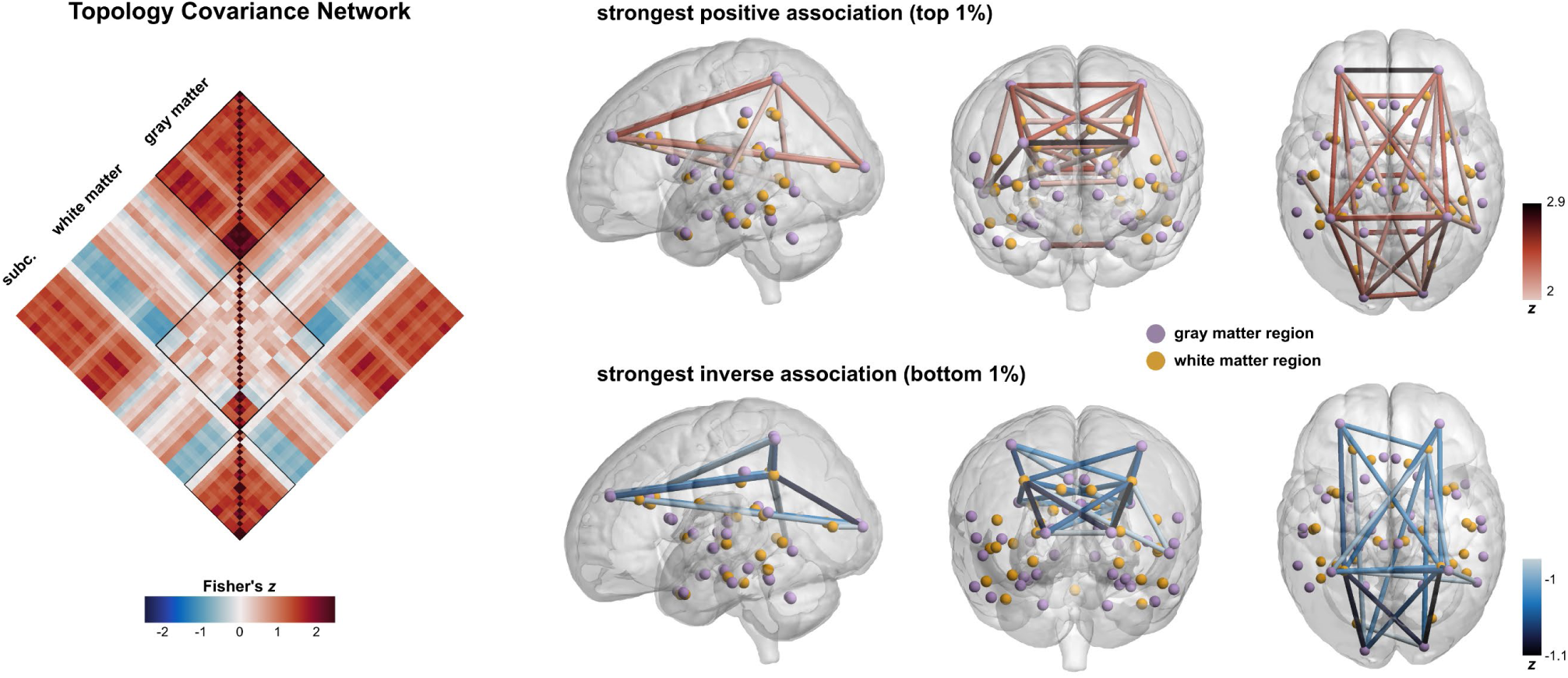
Covariance network of brain topology. The network displays the region-to-region covariance of topological complexity values across individual newborns (n=782, as in Fig. 2A). The left-hand side displays the direction of how brain regions covary each other, showing primarily positive associations within regions of the same tissue compartment as well as for cortical gray matter and subcortical areas, while white matter regions show widespread inverse associations to subcortical areas and cortical gray matter. The right-hand side shows this network in brain space, thresholded to the strongest 1% of positive and inverse associations, respectively. Here, the strongest positive associations (top) are observed between areas of the same tissue class and homologous areas, and the strongest inverse associations (bottom) between cortical gray matter and white matter areas.

**Figure S2.**
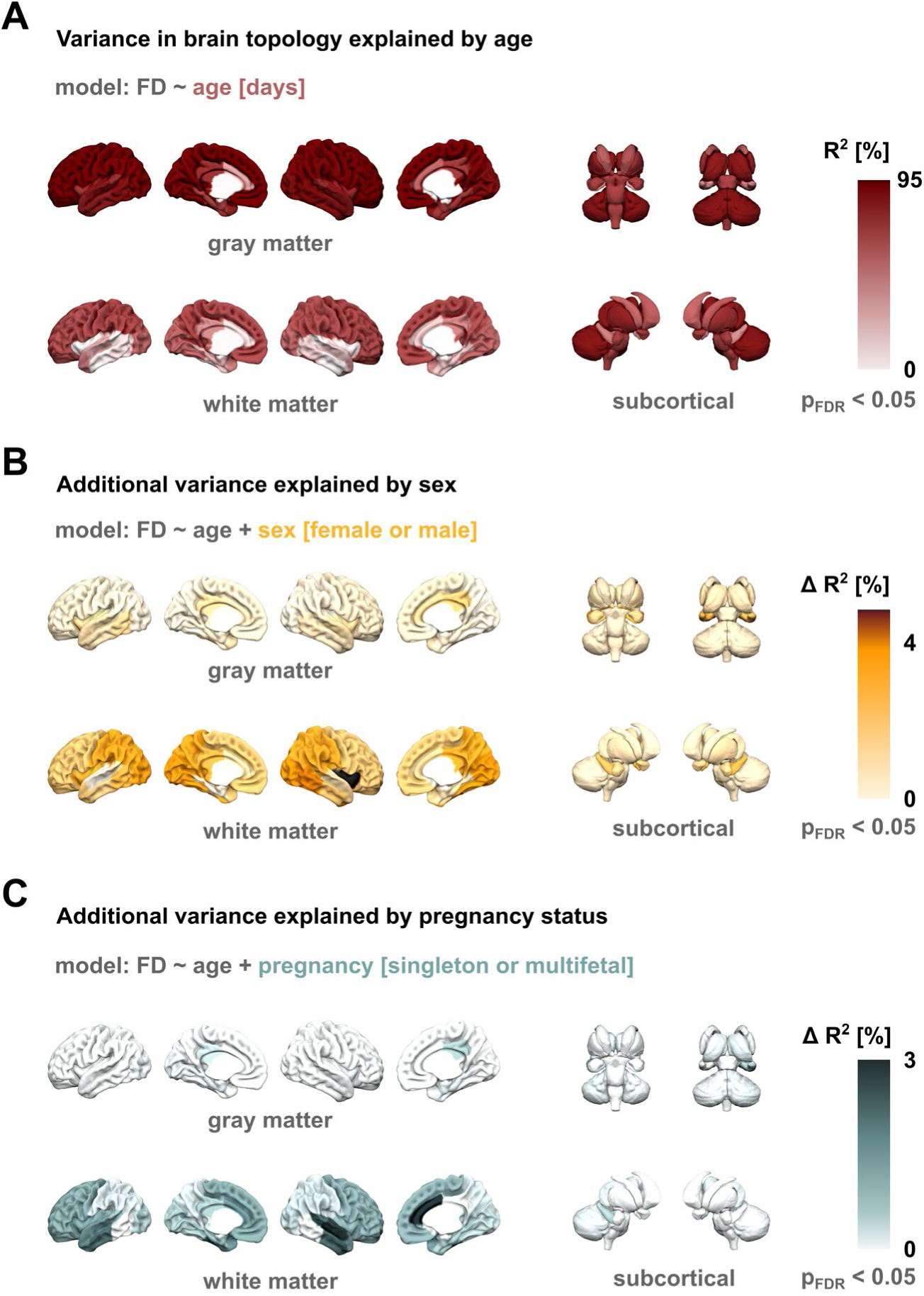
Explaining variance in brain shape with age, sex, and pregnancy status. **A**, Variance in fractal dimensionality explained by age at scan in a compact linear regression model. **B**, Hierarchical regression results showing the additional variance explained (ΔR^2^) by including the sex of the infant into the age model. **C**, Hierarchical regression results showing the additional variance explained by including the pregnancy status (singleton or multifetal) into the age model.

**Figure S3.**
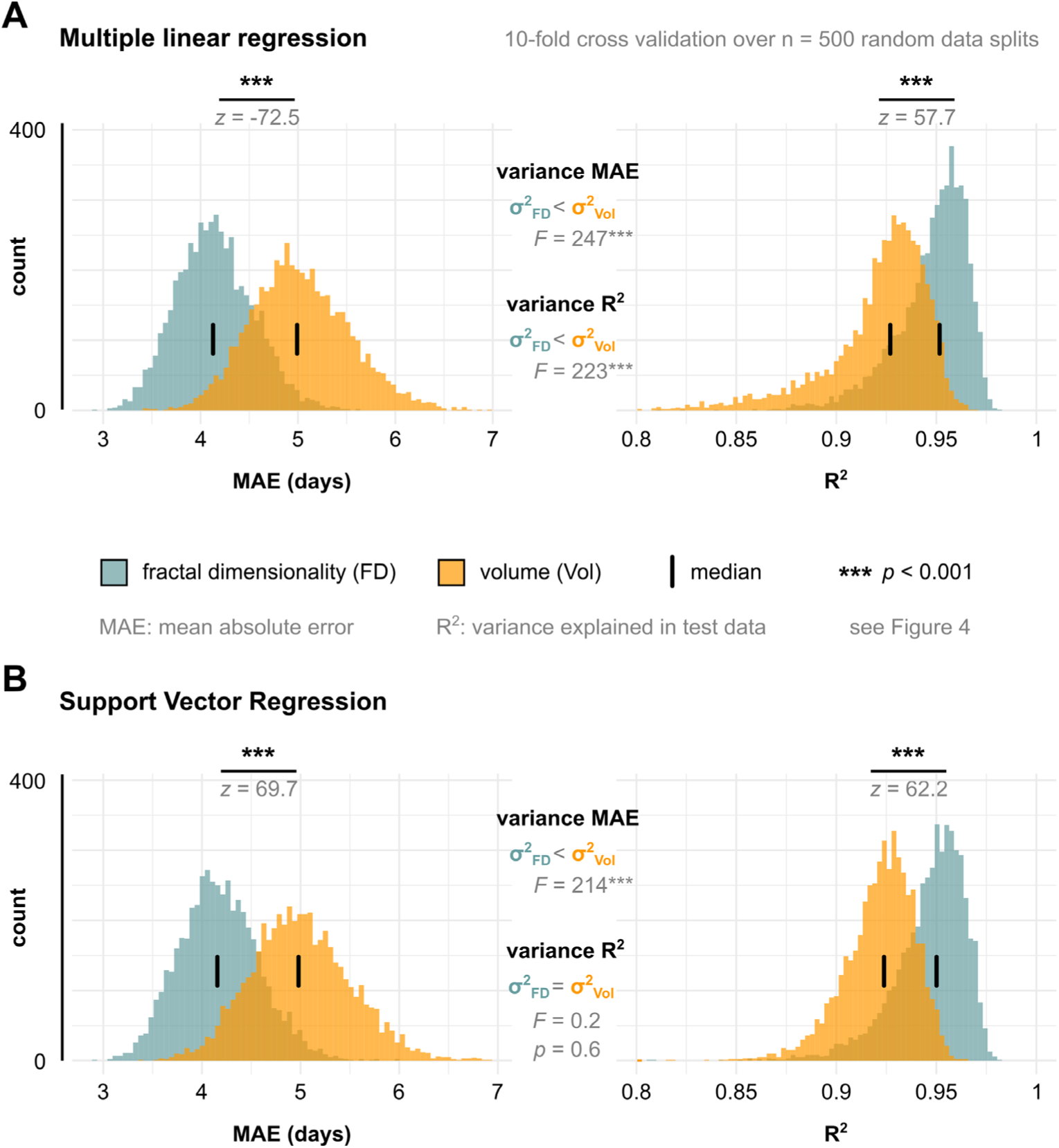
The superior performance of age prediction from fractal dimensionality over age prediction from volume is equivalently observed in two alternative control models. **A**, Results of running the age prediction pipeline outlined in Figure 4A with simple multiple linear regression. The distributions of the performance metrics (MAE: mean absolute prediction error in days; R^2^: variance explained in test data) are obtained from 500 random repetitions of the 10-fold cross validation procedure, as in Figure 4C. **B**, Running the age prediction pipeline with Support Vector Regression (linear kernel). Across both these simpler modelling approaches, shape-based age prediction with fractal dimensionality (FD) consistently outperforms size-based age prediction with volume (Vol).

**Figure S4.**
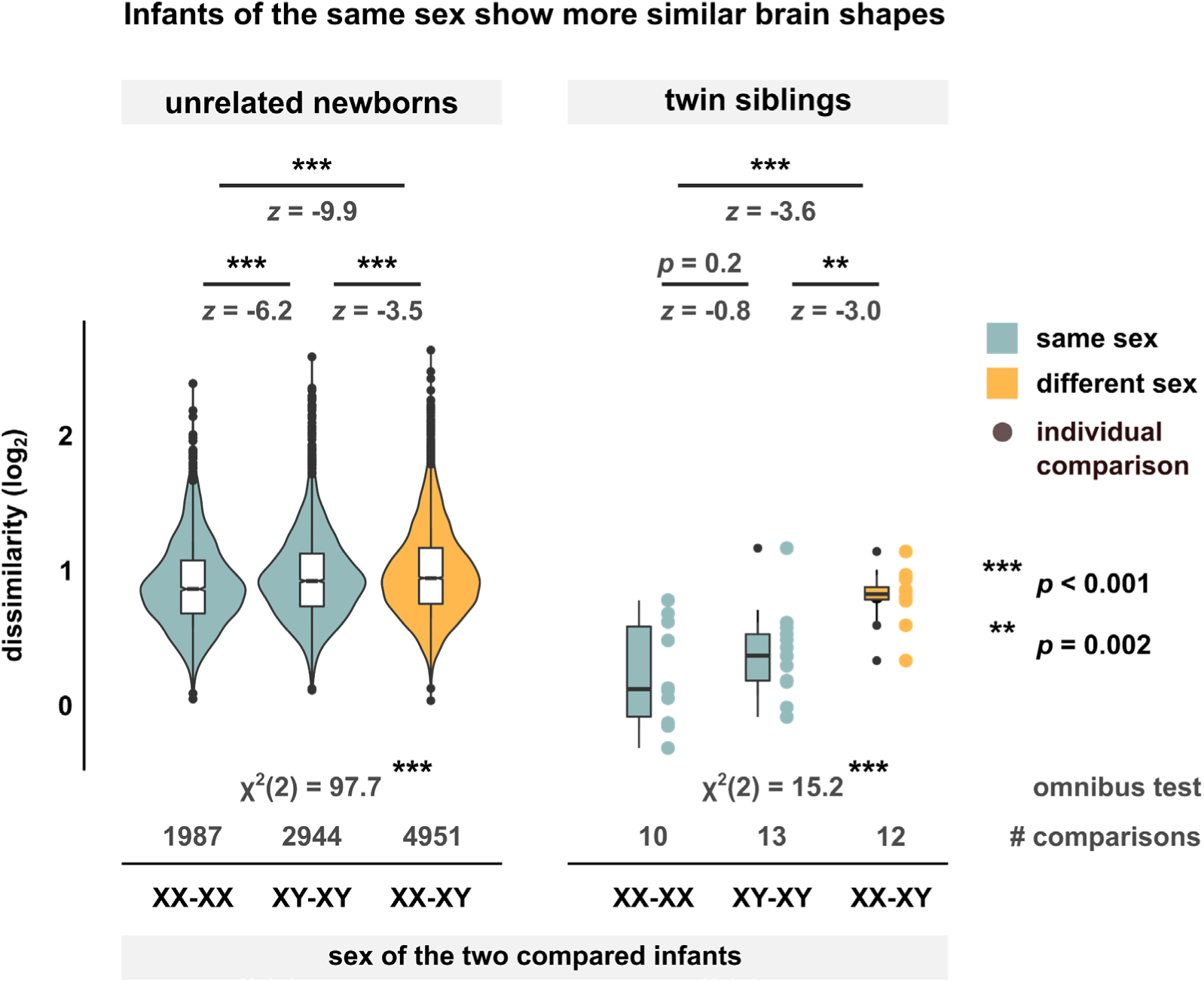
The shape similarity of any two brains depends on the sex of the compared infants. Dissimilarity scores of all comparisons between newborns that were within 1 day of age at the time of scanning (left: unrelated infants, right: twin siblings). Dissimilarity is displayed on a binary logarithmic scale and is stratified by the sex of the compared infants (XX-XX: both infants female; XY-XY: both infants male; XX-XY: one female, one male). Omnibus tests correspond to Kruskal-Wallis tests, and pairwise comparisons to Dunn’s tests with FDR adjustment.

**Figure S5.**
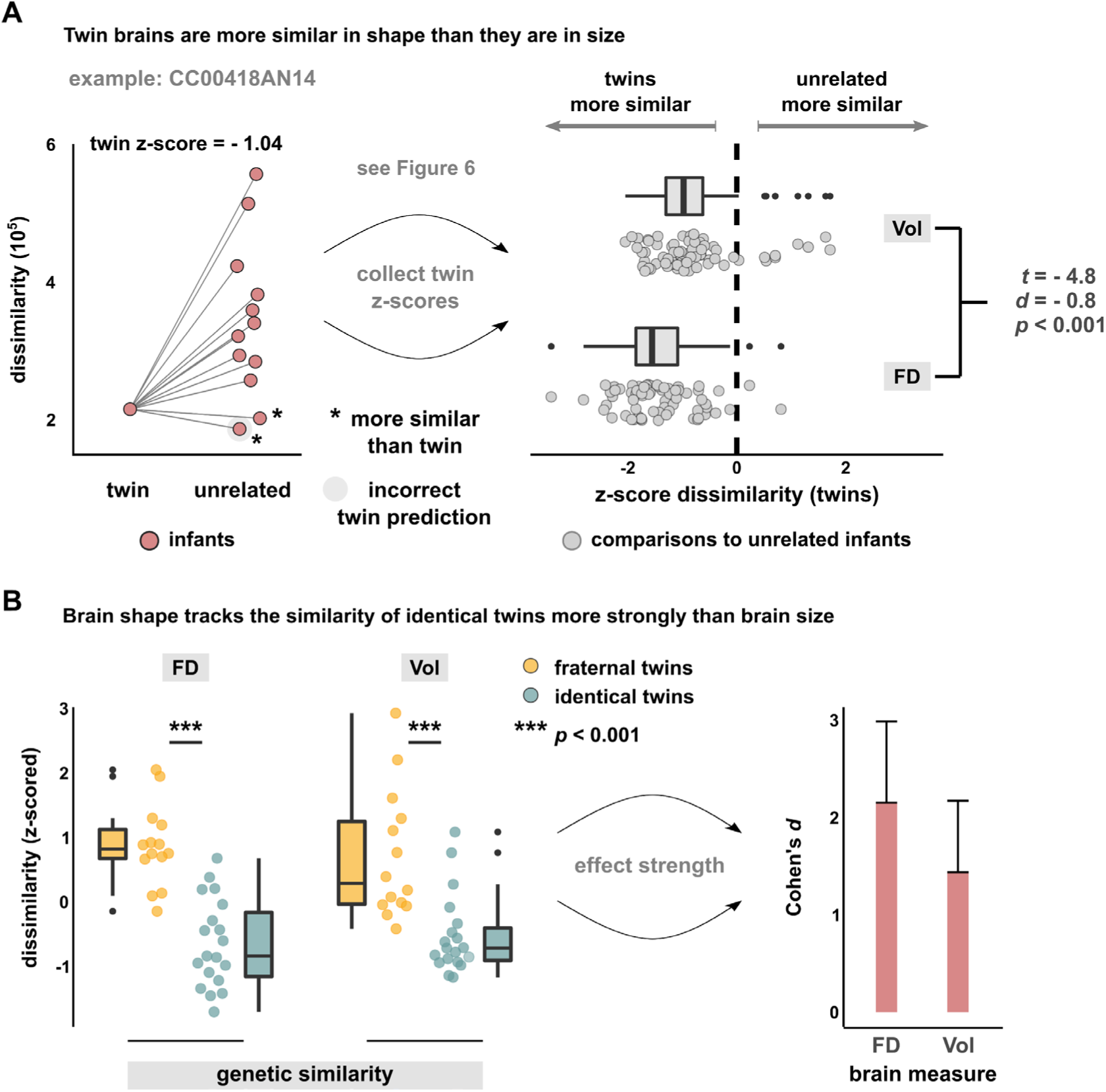
Brain size is less sensitive to genetic similarity than brain shape. **A**, Dissimilarity of brain volumes (Vol). Comparisons of the exemplary infant in main figure 6C to its twin sibling and the set of age-matched unrelated infants. Note that in contrast to the results with fractal dimensionality (FD), two unrelated newborns are more similar to the exemplary infant than its twin sibling in terms of brain size. Consequently, with volume, an unrelated infant is incorrectly predicted to be the twin of the exemplary infant, as twin prediction is based on the lowest-ranking dissimilarity score (see Fig. S6 for a comprehensive comparison of shape-based vs. size-based twin prediction). The right panel compares the dissimilarity scores for all twin-to-unrelated comparisons between volume and fractality (Welch’s two-sample t-test), where the latter corresponds to the distribution of main Figure 6D. **B**, Standardized dissimilarity scores for volume and fractal dimensionality. While both measures yield lower differences for identical twin pairs compared to fraternal twins, the strength of this effect is higher for fractal dimensionality than for volume (error bars: 95% confidence interval).

**Figure S6.**
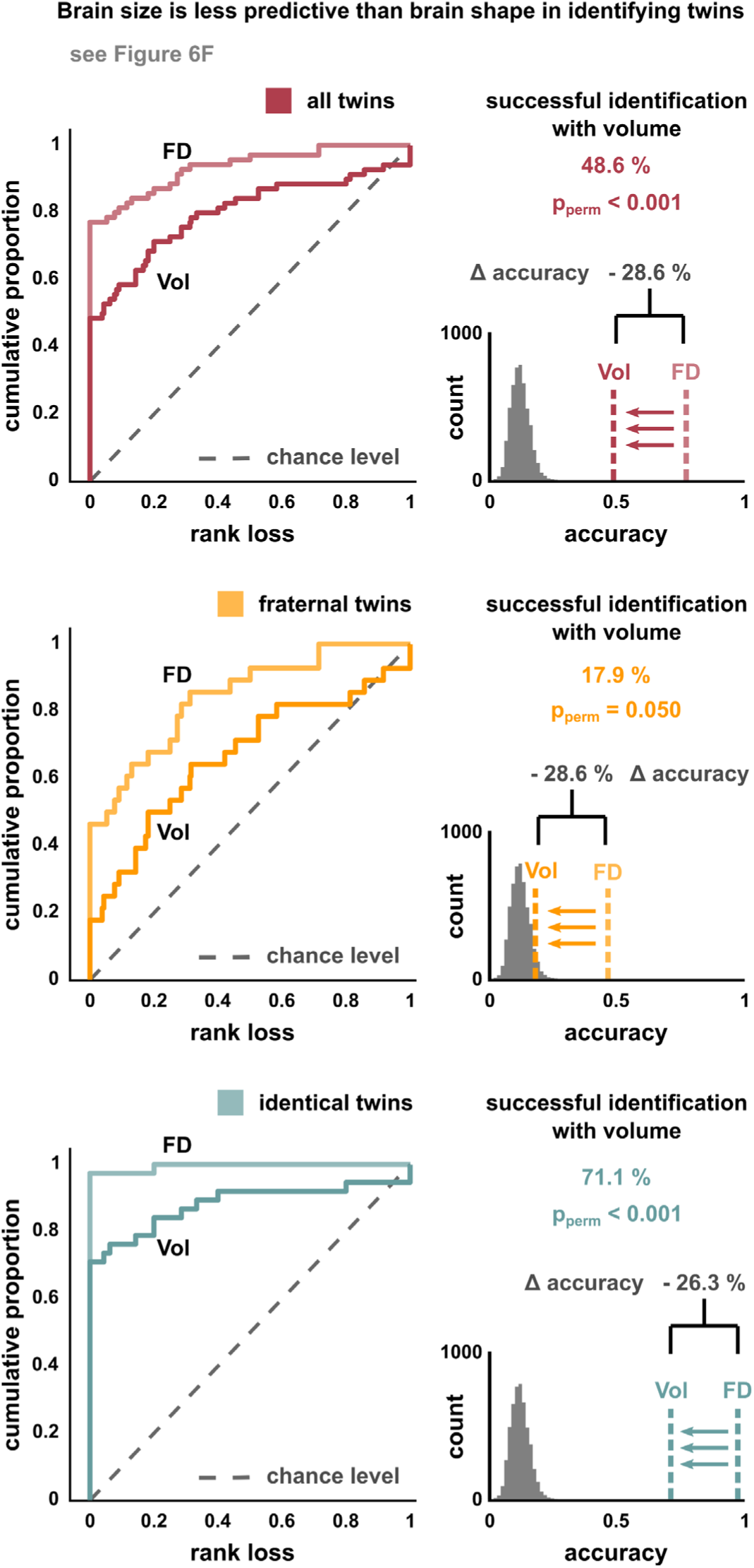
Brain size is less predictive than brain shape in identifying twin siblings. The figure relates the predictive capacity of identifying twin siblings with volume (Vol) instead of fractal dimensionality (FD). Compared to twin prediction from shape (see Fig. 6F in main text), twin prediction from brain size showed a consistent 25-30% drop in identification accuracy for all twin predictions (top row), the subgroup of fraternal twins (middle), and the subgroup of identical twins (bottom). Gray histograms correspond to the null distribution of correct twin identifications that happen by chance, as in Figure 6F.

